# COVID-ONE-humoral immune: The One-stop Database for COVID-19-specific Antibody Responses and Clinical Parameters

**DOI:** 10.1101/2021.07.29.454261

**Authors:** Zhaowei Xu, Yang Li, Qing Lei, Likun Huang, Dan-yun Lai, Shu-juan Guo, He-wei Jiang, Hongyan Hou, Yun-xiao Zheng, Xue-ning Wang, Jiaoxiang Wu, Ming-liang Ma, Bo Zhang, Hong Chen, Caizheng Yu, Jun-biao Xue, Hai-nan Zhang, Huan Qi, Siqi Yu, Mingxi Lin, Yandi Zhang, Xiaosong Lin, Zongjie Yao, Huiming Sheng, Ziyong Sun, Feng Wang, Xionglin Fan, Sheng-ce Tao

## Abstract

Coronavirus disease 2019 (COVID-19), which is caused by SARS-CoV-2, varies with regard to symptoms and mortality rates among populations. Humoral immunity plays critical roles in SARS-CoV-2 infection and recovery from COVID-19. However, differences in immune responses and clinical features among COVID-19 patients remain largely unknown. Here, we report a database for COVID-19-specific IgG/IgM immune responses and clinical parameters (COVID-ONE humoral immune). COVID-ONE humoral immunity is based on a dataset that contains the IgG/IgM responses to 21 of 28 known SARS-CoV-2 proteins and 197 spike protein peptides against 2,360 COVID-19 samples collected from 783 patients. In addition, 96 clinical parameters for the 2,360 samples and information for the 783 patients are integrated into the database. Furthermore, COVID-ONE humoral immune provides a dashboard for defining samples and a one-click analysis pipeline for a single group or paired groups. A set of samples of interest is easily defined by adjusting the scale bars of a variety of parameters. After the “START” button is clicked, one can readily obtain a comprehensive analysis report for further interpretation. COVID-ONE-humoral immune is freely available at www.COVID-ONE.cn.

## Introduction

COVID-19 is an unprecedented global threat caused by severe acute respiratory syndrome coronavirus 2 (SARS-CoV-2), which has already caused 188,843,580 infections and claimed 4,065,400 lives as of July 16, 2021 (https://coronavirus.jhu.edu/map.html) [1]. There is still no effective medicine [2, 3] for treating COVID-19.

Most patients recover via their own immunity, including SARS-CoV-2-specific IgG responses, especially neutralizing antibodies [4-6]. Overall, it is of great interest to decipher SARS-CoV-2-specific IgG and IgM responses at a systems level and to correlate responses to clinical parameters.

To understand how the human immune system responds to SARS-CoV-2, we constructed a SARS-CoV-2 proteome microarray containing 18 of the 28 predicted proteins and applied it to characterize IgG and IgM antibodies in the sera of 29 convalescent patients [7]. Recently, we upgraded the SARS-CoV-2 protein microarray, and the new microarray contains 21 predicted SARS-CoV-2 proteins and 197 spike protein peptides (with full coverage of spike) [8]. Using this microarray, we screened 2,360 serum samples from 783 COVID-19 patients, covering mild, severe and critical cases. Thus, we compiled a dataset with comprehensive information on SARS-CoV-2-specific humoral responses and rich in clinical parameters.

To share the dataset efficiently, in addition to the related research that we have already published [9-13], we built a database for COVID-19-specific humoral immune responses and clinical parameters, namely, COVID-ONE-humoral immune (www.covid-one.cn), using Shiny. This database contains a comprehensive dataset of IgG and IgM responses to the 21 predicted SARS-CoV-2 proteins and 197 spike protein peptides from a cohort of 783 COVID-19 patients. To bolster clinical relevance, 96 clinical parameters and basic patient information were also included. COVID-ONE humoral immunity provides search, data analysis, and visualization functions. In particular, COVID-ONE-humoral immune integrates antibody response landscape analysis, correlation analysis, machine learning, etc. In the data analysis module, users can easily define sample groups of interest by adjusting scale bars, and the sample groups can be either one group or paired groups. In-depth analysis is achieved by clicking a single button; optionally, the results can be saved and downloaded as an independent package for further analysis.

To our knowledge, COVID-ONE humoral immune is the first database for SARS-CoV-2-specific humoral immune responses. We believe that COVID-19 humoral immune will be of broad interest and will facilitate understanding of immune responses in COVID-19 to combat the pandemic.

## Materials and methods

### Patients and samples

All 783 COVID-19 cases were laboratory confirmed; the patients were hospitalized at Tongji Hospital from 25 January 2020 to 28 April 2020. The criteria for defining severity, *i*.*e*., mild, severe and critically severe, referenced the Diagnosis and Treatment Protocol for Novel Coronavirus Pneumonia (Trial Version 7), as released by the National Health Commission & State Administration of Traditional Chinese Medicine. For many of the patients, sera were collected during hospitalization at several time points. Negative reference samples were obtained from the National Institutes for Food and Drug Control. All serum samples were stored at -80°C until use.

### Peptide preparation

In this study, the SARS-CoV-2 spike protein (1,273 aa) was divided into 211 peptides of 12 aa, with 6 aa overlapping between adjacent peptides. After cysteine was added to the N-terminus, these peptides were synthesized by GL Biochem, Ltd. (Shanghai, China) and conjugated to BSA using Sulfo-SMCC (Thermo Fisher Scientific, MA, USA). Briefly, BSA was activated by Sulfo-SMCC at a molar ratio of 1:30 and dialyzed against PBS buffer. A total of 197 soluble peptides were individually conjugated with activated BSA in a w/w ratio of 1:1 and incubated for 2 h at room temperature. Free peptides were removed by dialysis with a pore size of 10 kD. The conjugates were assessed by SDS-PAGE.

### Protein preparation

SARS-CoV-2 protein sequences were downloaded from GenBank (Accession number: MN908947.3) and converted to *Escherichia coli* codon-optimized gene sequences. The optimized genes were synthesized and cloned into pET32a or pGEX-4T-1 by Sangon Biotech (Shanghai, China). Recombinant proteins were expressed in *E. coli* BL21 by growing cells in 200 mL LB medium to OD_600_=∼0.6 at 37 °C followed by induction with 0.2 mM isopropyl-β-d-thiogalactoside (IPTG) overnight at 16 °C. For the purification of 6xHis-tagged proteins, cell pellets were re-suspended in lysis buffer containing 50 mM Tris-HCl, 500 mM NaCl, and 20 mM imidazole (pH 8.0) and lysed using a high-pressure cell cracker (Union Biotech, Shanghai, China). After centrifugation at 12,000 x g for 20 min at 4 °C, the lysates were incubated with Ni^2+^ Sepharose beads (Senhui Microsphere Technology, Suzhou, China) for 1 h at 4 °C, washed 3 times with lysis buffer and eluted with buffer containing 50 mM Tris-HCl, 500 mM NaCl, and 300 mM imidazole (pH 8.0). For the purification of GST-tagged proteins, cells were harvested and lysed by a high-pressure cell cracker in lysis buffer containing 50 mM Tris-HCl, 500 mM NaCl, and 1 mM DTT at pH 8.0. After centrifugation, the supernatant was incubated with GST-Sepharose beads (Senhui Microsphere Technology, Suzhou, China). The target proteins were washed with lysis buffer and eluted with 50 mM Tris-HCl, 500 mM NaCl, 1 mM DTT, and 40 mM glutathione at pH 8.0. The purified proteins were quality checked by SDS-PAGE and Coomassie blue staining and stored at -80°C until use.

### Protein microarray fabrication

The SARS-CoV-2 proteome microarray used in this study is an updated version of the original microarray[7], which contains 18 of the 28 predicted SARS-CoV-2 proteins. Three more proteins, *i*.*e*., ORF3a, ORF3b, and ORF7b, and 197 spike protein peptides were added to the updated version. Therefore, the protein microarray used in this study contained 21/28 SARS-CoV-2 proteins and 197 peptides, with full coverage of the spike protein. The proteins and spike protein peptides, along with BSA and anti-human IgG/IgM (Jackson ImmunoResearch Laboratories, USA), were used as negative and positive controls, respectively, and printed in triplicate on PATH substrate slides (Grace Bio-Labs, Oregon, USA) to generate identical arrays in a 2 × 7 subarray format using a Super Marathon printer (Arrayjet, UK). Anti-His (Millipore, USA), anti-GST (Sigma, USA), and anti-BSA (Sangon Biotech, China) antibodies were used for quality control of the SARS-CoV-2 proteome microarray. The protein microarrays were stored at -80°C until use.

### Microarray-based serum analysis

A 14-chamber rubber gasket was mounted onto each slide to create individual chambers for 14 identical subarrays. The microarray was used for serum profiling as described previously, with minor modifications[14]. Briefly, arrays stored at -80°C were warmed to room temperature and then incubated in blocking buffer (3% BSA in 1×PBS buffer with 0.1% Tween 20) for 3 h. A total of 200 μL of diluted serum or antibodies was incubated with each subarray for 2 h. For most samples, sera were diluted to 1:200; for the competition experiment, free peptides were added at a concentration of 0.25 mg/mL. For the enriched antibodies, 0.1-0.5 μg antibodies were included in 200 μL incubation buffer. The arrays were washed with 1× PBST, and the bound antibodies were monitored by incubating with Cy3-conjugated goat anti-human IgG and Alexa Fluor 647-conjugated donkey anti-human IgM (Jackson ImmunoResearch, PA, USA) diluted 1:1,000 in 1× PBST at room temperature for 1 h. The microarrays were then washed with 1×PBST, dried by centrifugation at room temperature and scanned using a LuxScan 10K-A (CapitalBio Corporation, Beijing, China) with the parameters set as 95% laser power/PMT 550 and 95% laser power/PMT 480 for IgM and IgG, respectively. The fluorescence intensity was extracted with GenePix Pro 6.0 software (Molecular Devices, CA, USA).

### Protein microarray data analysis

IgG and IgM signal intensities were defined as foreground medians (F) subtracted by background medians (B) for each spot, and the signal intensity of a protein was averaged for triplicate spots. Block **#**14 of each slide was incubated with SARS-CoV-2 immunopositive serum as the positive control. Data normalization between slides was performed by a linear method according to the positive control; specifically, a normalization factor for each slide was calculated by linear regression according to the positive control. To reduce error among microarrays, the signals of all the proteins from each slide were divided by its normalization factor.

### Quantification and statistical analysis

To calculate the rate of antibody response for each protein, the mean plus 2 times the standard deviation (SD) of the control serum was set as the cut-off. R was used for most data analysis and drawing, *i*.*e*., Pearson correlation coefficient, ROC, T-test, cluster analysis and machine learning.

### Data collection

Specific IgG/IgM immune response data were obtained by microarray-based serum analysis. Blood parameters were collected from Tongji Hospital, Tongji Medical College, Huazhong University of Science and Technology, Wuhan, China.

### Database architecture and web interface

COVID-ONE-humoral immune is a Shiny-based (1.5.0) database. Shinydashboard (0.7.1) and Shiny BS (0.61) were used to shape the UI, and the package DT (0.15) was used to format data tables. For data analysis, dplyr (1.0.2), tidyverse (1.3.0), randomForest (4.6-14), pROC (1.16.2), and umap (0.2.6.0) were integrated into Shiny. Pheatmap (1.0.12) and ggplot2 (3.3.2) carry out plotting. For the basic environment, the operation system is Ubuntu 20.04 LTS, and the version of R is 3.6.3.

### Ethics statement

The study was approved by the Ethical Committee of Tongji Hospital, Tongji Medical College, Huazhong University of Science and Technology, Wuhan, China (ITJ-C20200128). Written informed consent was obtained from all participants enrolled in this study.

## Results

### The database framework and clinical information for the patients

In this study, we collected 2,360 serum samples from 783 patients with an average age of 61.4 years and average onset time of 50 days. There were 387 males and 396 females and 369 non-severe, 309 severe, 105 critical cases. Regarding outcome, there were 723 survivors and 60 deaths **(Fig. 1 A, Table 1, Supplementary dataset 1)**.

**Figure 1.**
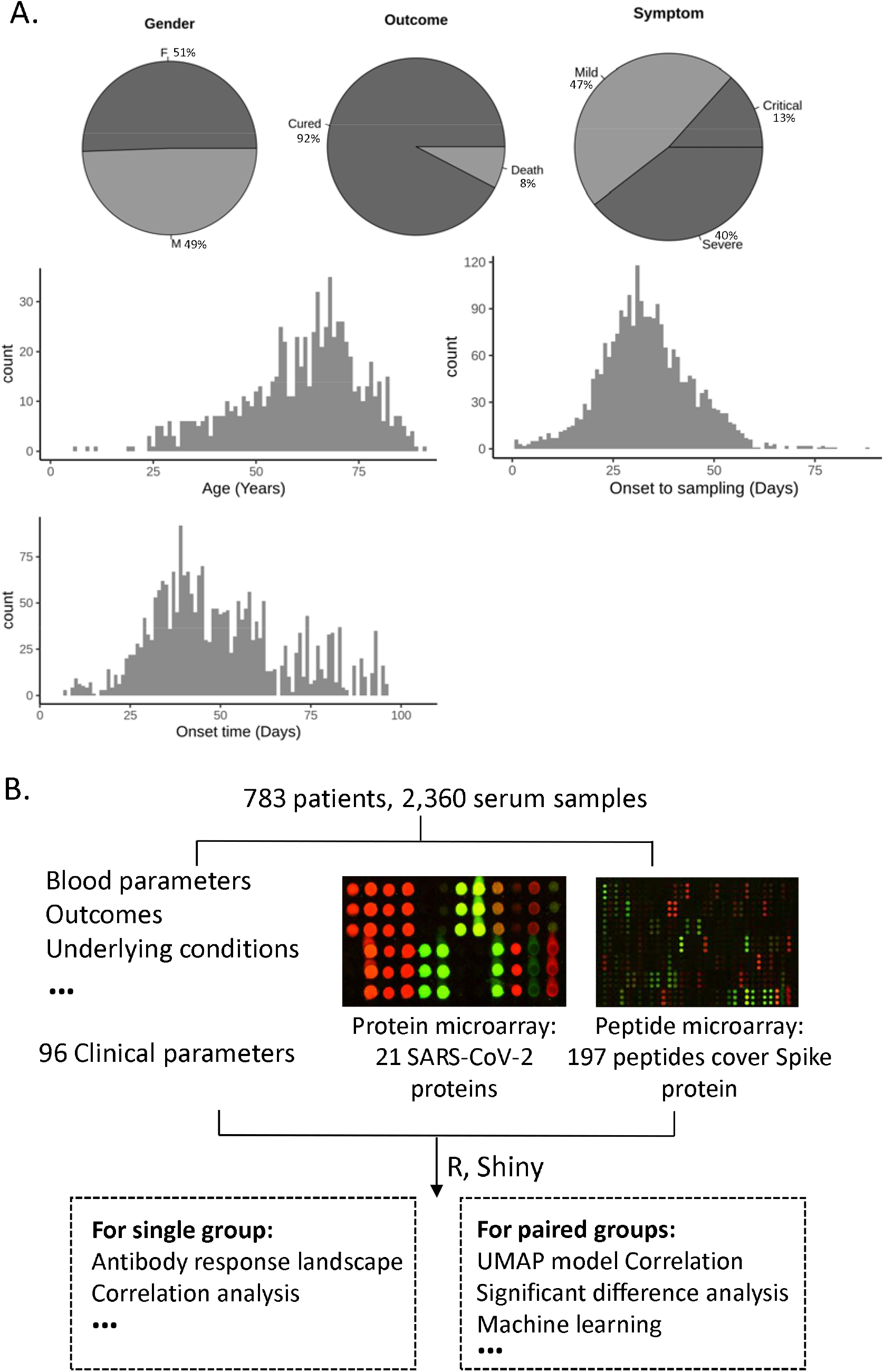
Overview of data resources and functional modules of COVID-ONE humoral immunity. **(A)** The patient information of the study cohort showing sex, outcome, severe type, *etc*. **(B)** The framework of COVID-ONE-humoral immune. The one-stop database for COVID-19-specific humoral immune responses and clinical parameters. The COVID-ONE humoral immune dataset includes 220 protein/peptide antibody responses and 96 clinical parameters from 2360 serum samples. Using the Shiny package, COVID-ONE-humoral immune provides single-group or paired-group analysis based on the dataset.

To systematically analyse immune responses to SARS-CoV-2 infection, we screened 2,360 serum samples using a COVID-19 protein microarray that contains 21 proteins and 197 spike protein peptides. Additionally, we analysed 89 blood parameters for the 2,360 serum samples, *i*.*e*., complete blood count, blood chemistry study and blood enzyme tests. Hence, we obtained a comprehensive dataset that contains SARS-CoV-2-specific humoral responses and is rich in clinical parameters.

By combining clinical information, IgG/IgM immune responses and blood parameters, we established a database (COVID-ONE humoral immune) that provides a one-stop analysis pipeline for COVID-19-specific immune responses and clinical parameters **(Fig. 1 B)**. To allow users to obtain more COVID-19 serum profiling data, we set up a page on the COVID-ONE humoral immune website, named “More studies”, to archive other highly related data of COVID-19 serum profiling (protein/peptide microarrays/phage display) [15-20]. In addition, a healthy control dataset was added to the HELP page, which contains the IgG and IgM responses for 528 healthy people to the 21 proteins and spike protein peptides (Supplementary dataset 2).

The following steps are included in the analysis module:

- Users select a set of samples in the panel of patient information and click START.
- COVID-ONE humoral immune filters candidate samples according to the given parameters.
- COVID-ONE-humoral immune conducts analysis and provides results on the webpage.

To demonstrate how to use COVID-ONE humoral immunity for analysis, we provide 2 datasets for a single group and paired groups as examples.

### Case□: Antibody responses and clinical parameters of non-survivors of COVID-19

To study features of COVID-19 non-survivors, we selected the “death” parameter of outcome in a single-group analysis module. This cohort contained 392 serum samples and 60 cases, with an average age of 69.6 years and sex (38 male, 22 female) **(Table 2)**.

The IgG response landscape analysis of SARS-CoV-2 proteins showed positive rates for the S and N proteins and ORF3b of 95%, 93% and 87%, respectively, consistent with previous studies [21, 22] **(Fig. 2A)**. Interestingly, non-structural protein 7 (NSP7) had an 88% IgG positive rate, which suggests that NSP7 may play an important role in COVID-19 **(Fig. 2A)**. Spike peptide S1-45 had the highest positive rate (87%) for the IgM response, indicating that the region including S1-45 may play an important role in IgM immunity **(Fig. S1)**.

**Figure 2.**
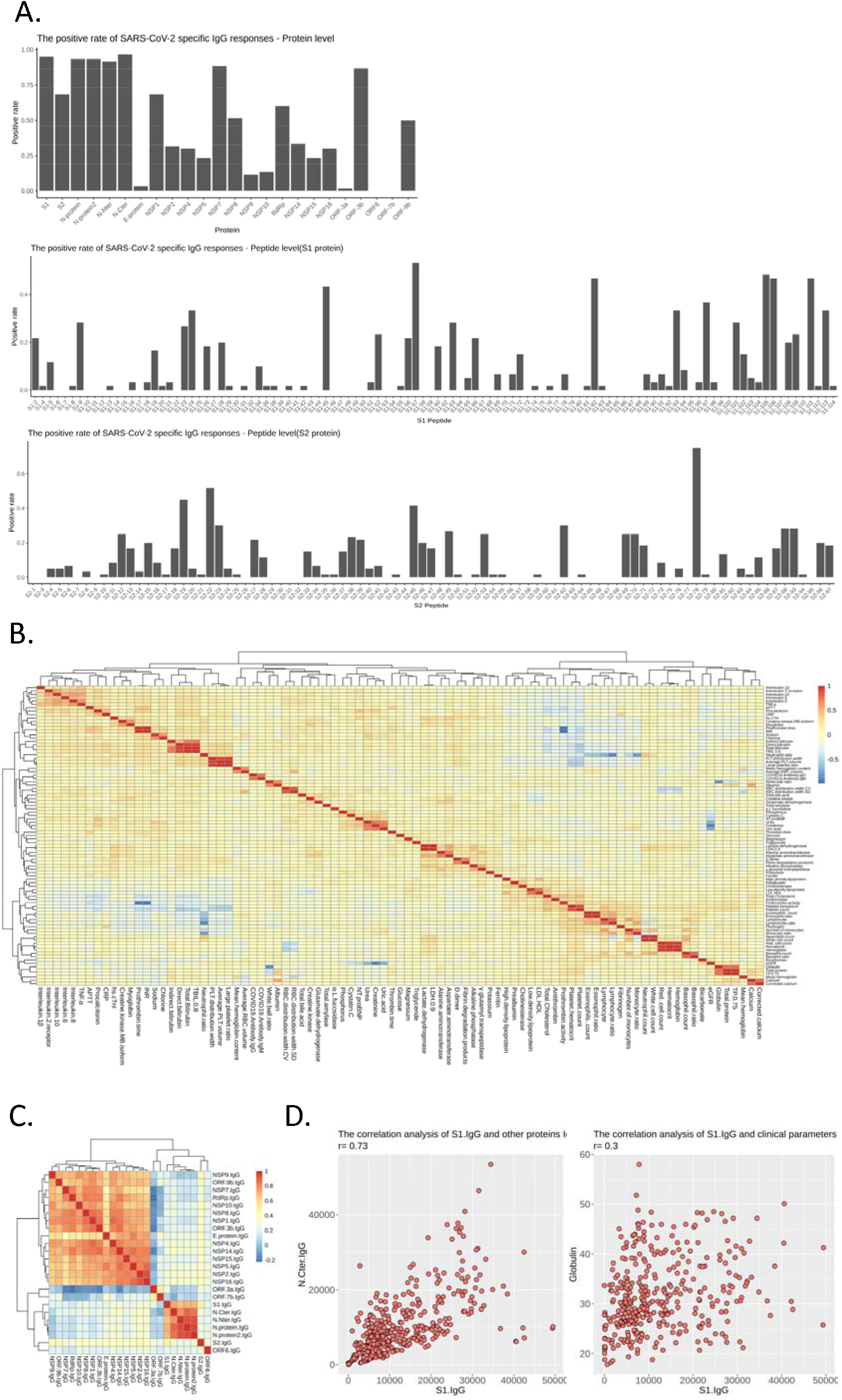
SARS-CoV-2-specific antibody responses and their correlations with clinical parameters: COVID-19 non-survivors. **(A)** The antibody IgG response landscape against SARS-CoV-2 proteins (upper part), S1 protein peptides (middle part) and S2 protein peptides (lower part). **(B)** Heat map showing correlation analysis of blood parameters. **(C)** Heat map showing correlation analysis of antibody IgG responses against SARS-CoV-2 proteins. **(D)** Scatter plot showing correlation analysis between the S1 IgG response and protein IgG responses/blood parameters.

Correlation analysis of clinical parameters showed that the neutrophil count had a negative correlation with the monocyte count and lymphocyte ratio **(Fig. 2B)**. In addition, correlation analysis of antibody IgG responses showed a high correlation for IgG responses of the S1 and N proteins, but not for S2, with all non-structural protein IgG responses having no or very weak correlations **(Fig. 2C)**. To study influencing factors of S1 antibody production, we analysed correlation between the S1 IgG response and clinical parameters and found the response to correlate with globulin in patients with critical COVID-19 **(Fig. 2D)**.

### Case□: Differences in IgG/IgM immune responses and clinical parameters associated with sex

Previous studies have shown that sex has a considerable effect on the outcome of COVID-19 [23, 24] and is associated with underlying differences in immune responses to infection [25]. To study differences in IgG/IgM immune responses and clinical parameters between the sexes, we defined Group A as female and Group B as male for severe patients, with 231 males at an average age of 64.3 and 183 females at an average age of 68.1. Consistent with previous studies [26], males had a higher risk of severe disease than females (231/377 vs 183/379, *p*<0.001) **(Table 3, Table 4)**.

UMAP results showed no overall difference in IgG immunity between males and females **(Fig. 3A)**. To explore the disease mechanism in the sexes, we performed in-depth analysis for antibody response and blood parameters using COVID-ONE. The antibody response landscape shows that male patients have a higher positive rate than females for ORF-9b IgG, RdRp IgG, NSP1 IgG, *etc*. **(Fig. 3B)**. Moreover, longitudinal antibody dynamic analysis showed a stronger ORF-9b IgG response in males during the whole period of symptom onset, with a stronger NSP1 IgG response during the early stage of symptom onset; however, there was no significant difference for RdRp IgG **(Fig. 3C-E)**. ORF-9b has been considered a drug target for the treatment of COVID-19 because it suppresses type I interferon responses[27-29]. To explore the relevance between ORF-9b antibody responses and COVID-19 severity, we compared ORF-9b antibody responses between mild and severe cases, and the results showed that males with severe disease had higher ORF-9b antibody responses than females **(Fig. 3G-H)**.

**Figure 3.**
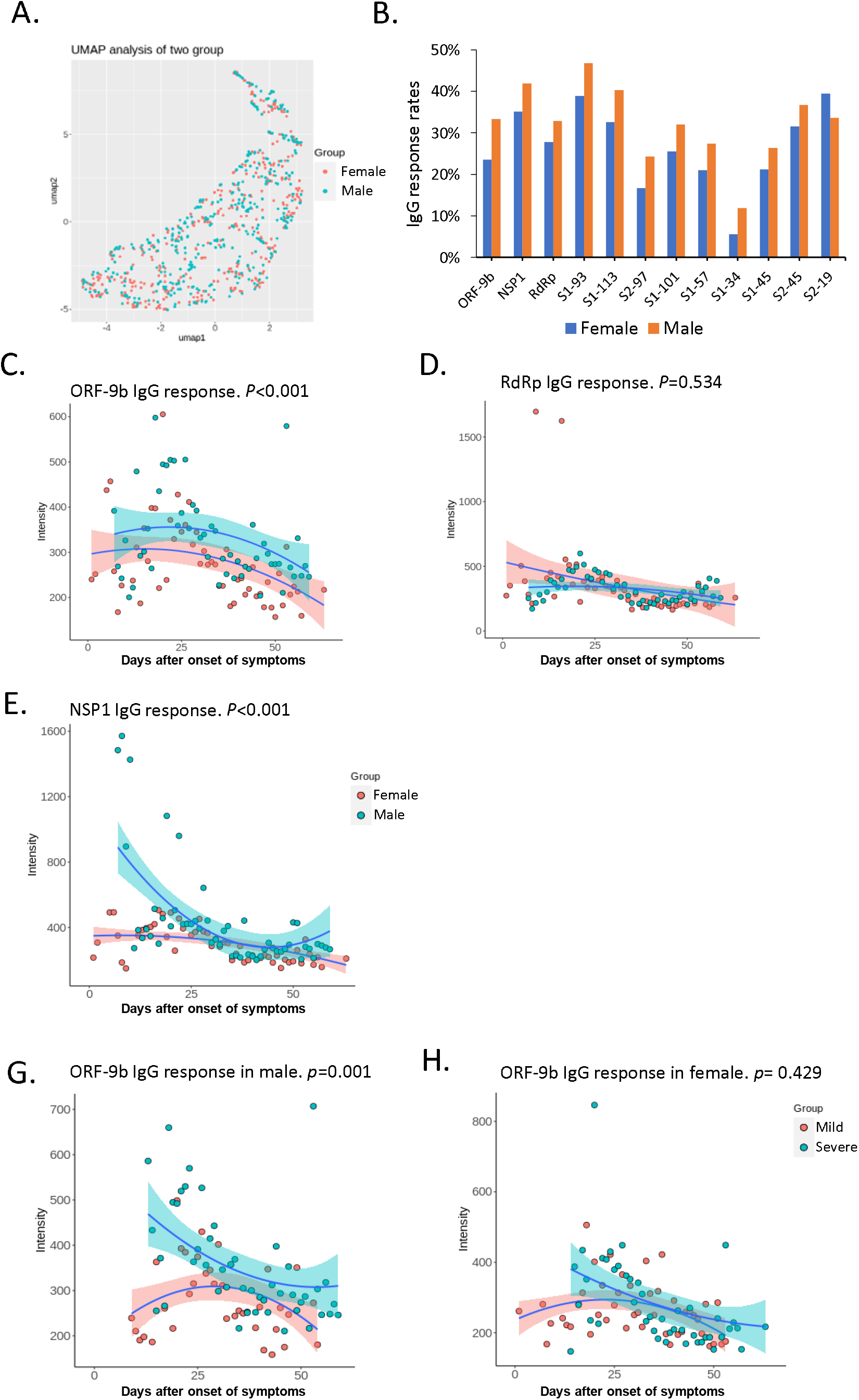
Correlation of the ORF-9b IgG response based on COVID-19 severity in male patients. **(A)** Scatter plot showing uniform manifold approximation and projection (UMAP) for serum samples using 21 protein IgG/IgM responses in sex subgroup analysis. **(B)** Histogram showing different responses in males and females for the IgG response. **(C-E)** Scatter plot showing ORF9b, RdRp and NSP1 IgG dynamic responses using longitudinal samples from male and female patients. **(G-H)** Scatter plot of the dynamic anti-ORF9b IgG response in COVID-19 patients with mild and severe symptoms.

To further decipher differences between female and male patients with COVID-19, we employed random forest for machine learning. The results showed creatinine, which is an acute kidney injury marker, to be the most significant factor between males and females **(Fig. 4A)**. To explore the relevance between creatinine and sex in COVID-19, we compared the level and dynamic response of creatinine in males and females and observed that the creatinine level in males was significantly higher than that in females **(Fig. 4B-C)**. To explore the relevance between creatinine and COVID-19 severity, we compared creatinine levels in mild and severe cases, and similar to ORF-9b antibody responses, male patients with severe COVID-19 had a higher level of creatinine **(Fig. 4D-E)**. Hence, ORF-9b antibodies and creatinine are associated with severe disease in male patients, which suggests different pathogeneses and complications between male and female COVID-19 patients.

**Figure 4.**
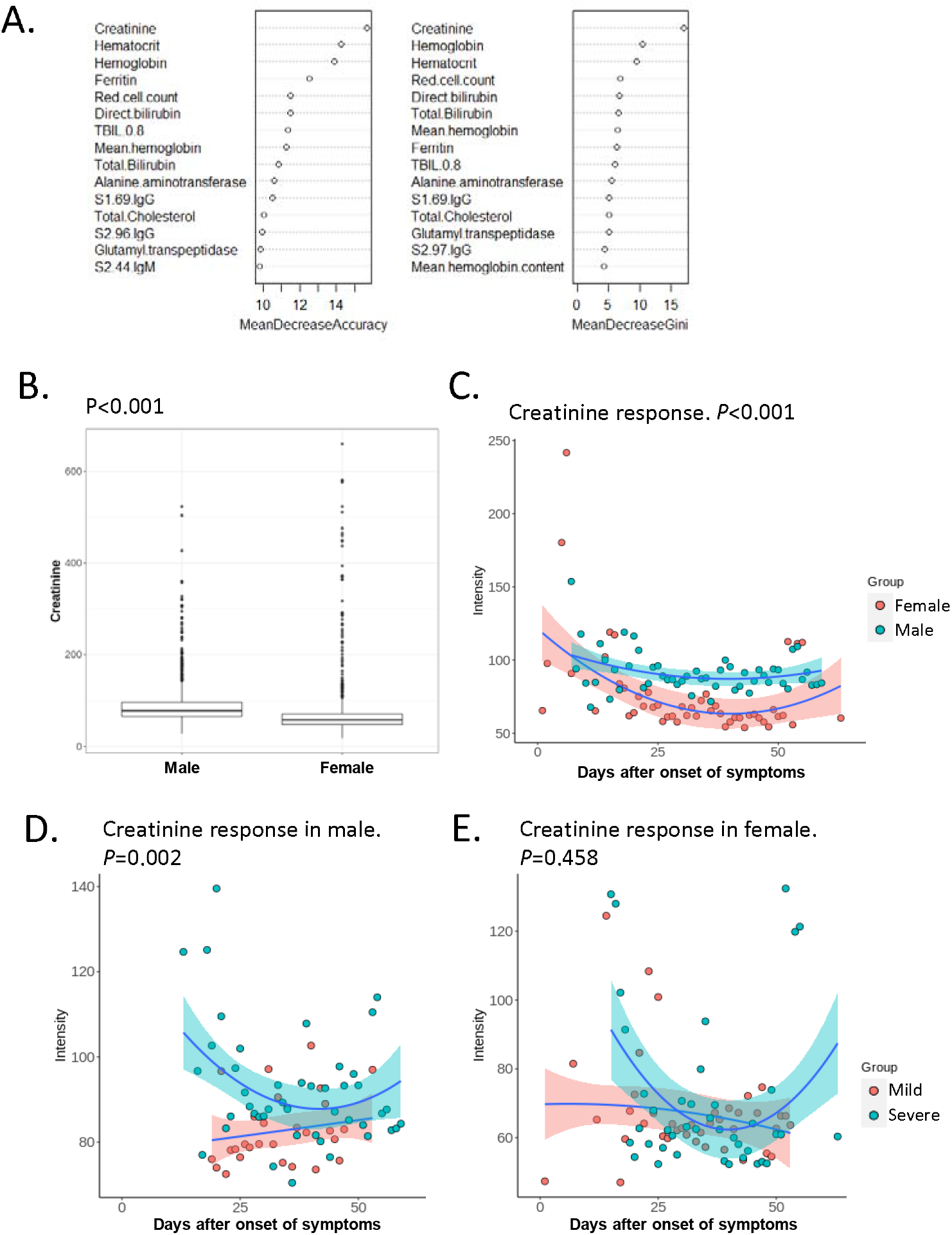
Correlation of creatinine response based on COVID-19 severity in male patients. **(A)** The top 15 sex-specific parameters by random forest analysis ranked by the mean decrease in accuracy and mean decrease in the Gini coefficient **(B)** The boxplot shows the significant difference in creatinine in sex subgroup analysis. The P-value was calculated by a two-sided t-test. **(C)** Scatter plot of creatinine levels of male and female COVID-19 patients. **(D-E)** Scatter plot of creatinine levels of COVID-19 patients with mild and severe disease.

## Discussion

In this study, we built COVID-ONE humoral immune, a COVID-19-specific database, using R Shiny. COVID-ONE humoral immune is based on a comprehensive dataset generated by analysing 2,360 COVID-19 sera using a SARS-CoV-2 protein microarray containing 21 of the 28 known SARS-CoV-2 proteins and 197 peptides completely covering the entire S protein sequence.

There are several published studies identifying the clinical characteristics, biomarkers and specific antibody responses of diverse COVID-19 patients **(Table S1)**. To strengthen the credibility of our dataset, we compared COVID-19-specific antibody responses with other studies at different levels. At the protein level, we analysed the dynamic responses to the S and N proteins. The results showed that S and N responses peaked at 6 weeks after the onset of symptoms for IgG and 4 weeks for IgM, which is consistent with the results of previous studies[19, 21] **(Fig. S2)**. At the peptide level, we compared IgG recognition of immunodominant regions in the SARS-CoV-2 spike protein and found that some high response areas that we identified[12] were consistent with those of Shrock et al. [15]: 25-36 aa, 553-588 aa, 770-829 aa, 1148-1159 aa and 1256-1273 aa. Another hot spot (aa 451-474) was only detected in our study. Regarding antibody diagnosis, Assia *et al*. achieved an AUC of 0.986 for IgG and 0.988 for IgM for the detection of prior SARS-CoV-2 infection when combining N and spike[20]. In our study, the AUC of the N protein was 0.995 for IgG and 0.988 for IgM, and the AUC of the S1 protein was 0.992 for IgG and 0.992 for IgM. We also found that S2–78 (1148–1159 aa) IgG is comparable to S1 IgG for COVID-19 patients, with an AUC of 0.99 for IgG and 0.953 for IgM[11].

To our knowledge, COVID-19 humoral immune is the first database for COVID-19-specific immune responses enriched in clinical parameters and has the following features. (i) Universality: COVID-ONE humoral immune contains 783 patients with 16 medical histories, which will be of broad interest for researchers and clinicians from diverse backgrounds. (ii) Accessibility: COVID-ONE-humoral immune provides a one-stop analysis pipeline, by which users can easily obtain meaningful information. (iii) Scalability: COVID-ONE humoral immune is built on the R platform, which is freely accessible, and many modular tools are readily available; thus, we can easily expand and incorporate new analyses for the dataset whenever necessary without changing the overall structure of the database. Nonetheless, there are some limitations for COVID-ONE humoral immunity. For example, it lacks data for convalescent patients, peptide-level humoral responses to proteins other than S protein, and multicentre samples. In the future, we will assay the dynamic responses of SARS-CoV-2-specific antibodies using ∼500 serum samples from ∼100 COVID-19 convalescent patients. We will also integrate published peptide microarray/phage display-related data[15-17, 30] and attempt to update the database covering the whole SARS-CoV-2 proteome at the peptide or amino acid level. In addition, the SARS-CoV-2 protein microarray has already been promoted by CDI Labs (www.cdi.bio) and ArrayJet (www.arrayjet.co.uk), and we anticipate more diverse data for SARS-CoV-2-specific antibody responses from multicentre samples. We strongly believe that by sharing a large dataset and facilitating data analysis, COVID-19 humoral immune is a valuable resource for COVID-19 research.

## Data and tool availability

COVID-ONE-humoral immune is freely accessible at www.covid-one.cn. The SARS-CoV-2 proteome microarray data are deposited on Protein Microarray Database under the accession number PMDE244 (http://www.proteinmicroarray.cn). If author need the raw data of antibody responses or clinical parameters, please contact the corresponding author.

## Author’s contributions

SCT and XLF developed the conceptual ideas and designed the study. ZWX, LKH, YL, QL, DYL, SJG, HWJ, HNZ, HQ, XL, performed the experiments and data analysis. ZWX, LKH, YL, XL built the database. SCT and ZWX wrote the manuscript with suggestions from other authors.

## Acknowledgments

This work was partially supported by the National Key Research and Development Program of China Grant (No.2016YFA0500600), National Natural Science Foundation of China (No. 31970130, 31600672, 31900112, 21907065 and 32000027).

## Figure legends

**Figure S1. The antibody IgM response landscape against SARS-CoV-2 proteins (upper part), S1 protein peptides (middle part) and S2 protein peptides (lower part)**.

**Figure S2. Dynamic antibody responses to S1 and N proteins**. Scatter plot showing dynamic antibody responses to S1 IgG **(A)**, N protein IgG **(B)**, S1 IgM **(C)**, and N protein IgM **(D)**.

