## Supplementary material for "COVID-ONE-humoral immune: The One-stop Database for COVID-19-specific Antibody Responses and Clinical Parameters": Table

### Slide 1
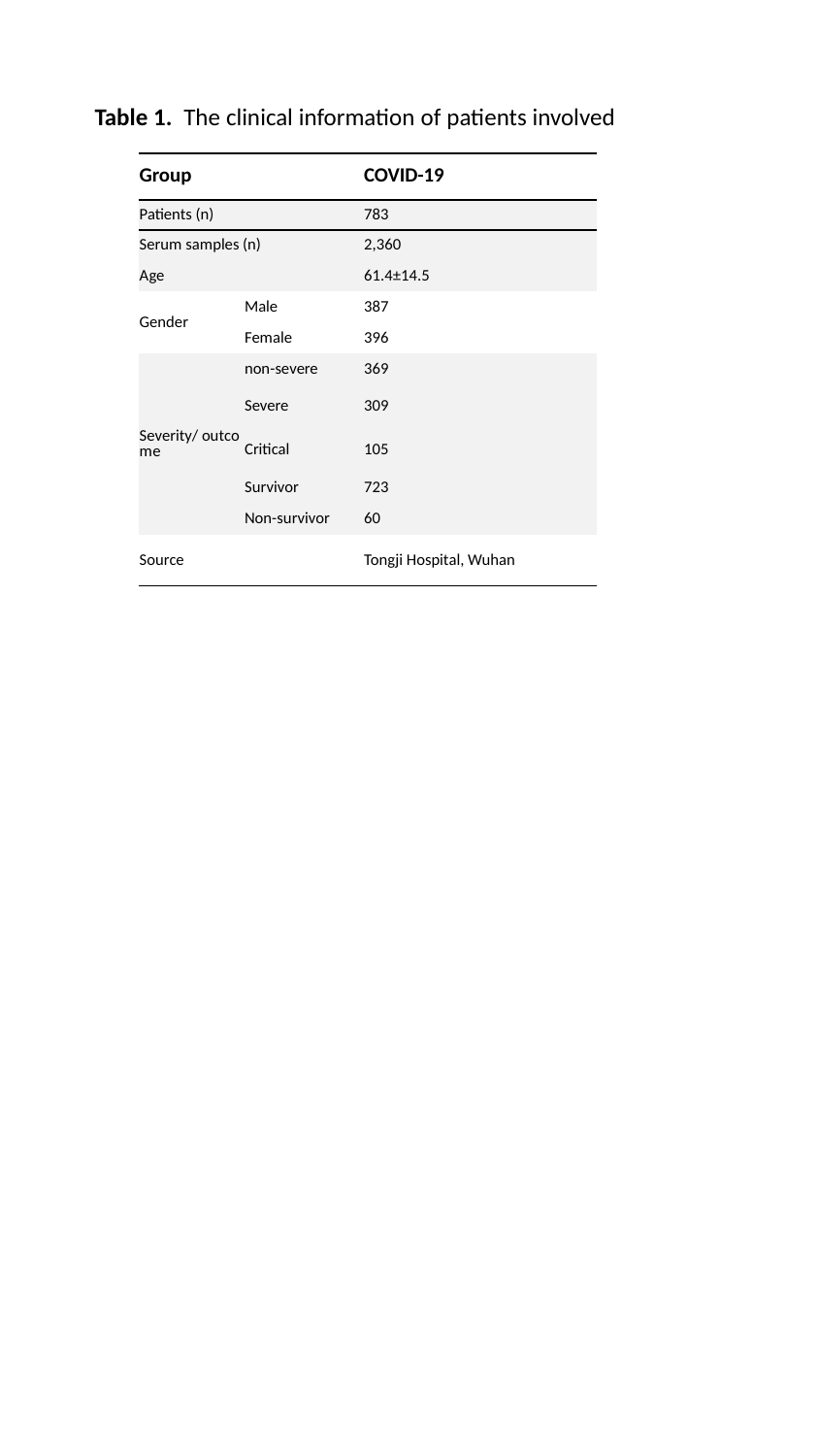

Table 1. The clinical information of patients involved
| Group | | COVID-19 |
| --- | --- | --- |
| Patients (n) | | 783 |
| Serum samples (n) | | 2,360 |
| Age | | 61.4±14.5 |
| Gender | Male | 387 |
| | Female | 396 |
| Severity/ outcome | non-severe | 369 |
| | Severe | 309 |
| | Critical | 105 |
| | Survivor | 723 |
| | Non-survivor | 60 |
| Source | | Tongji Hospital, Wuhan |

### Slide 2
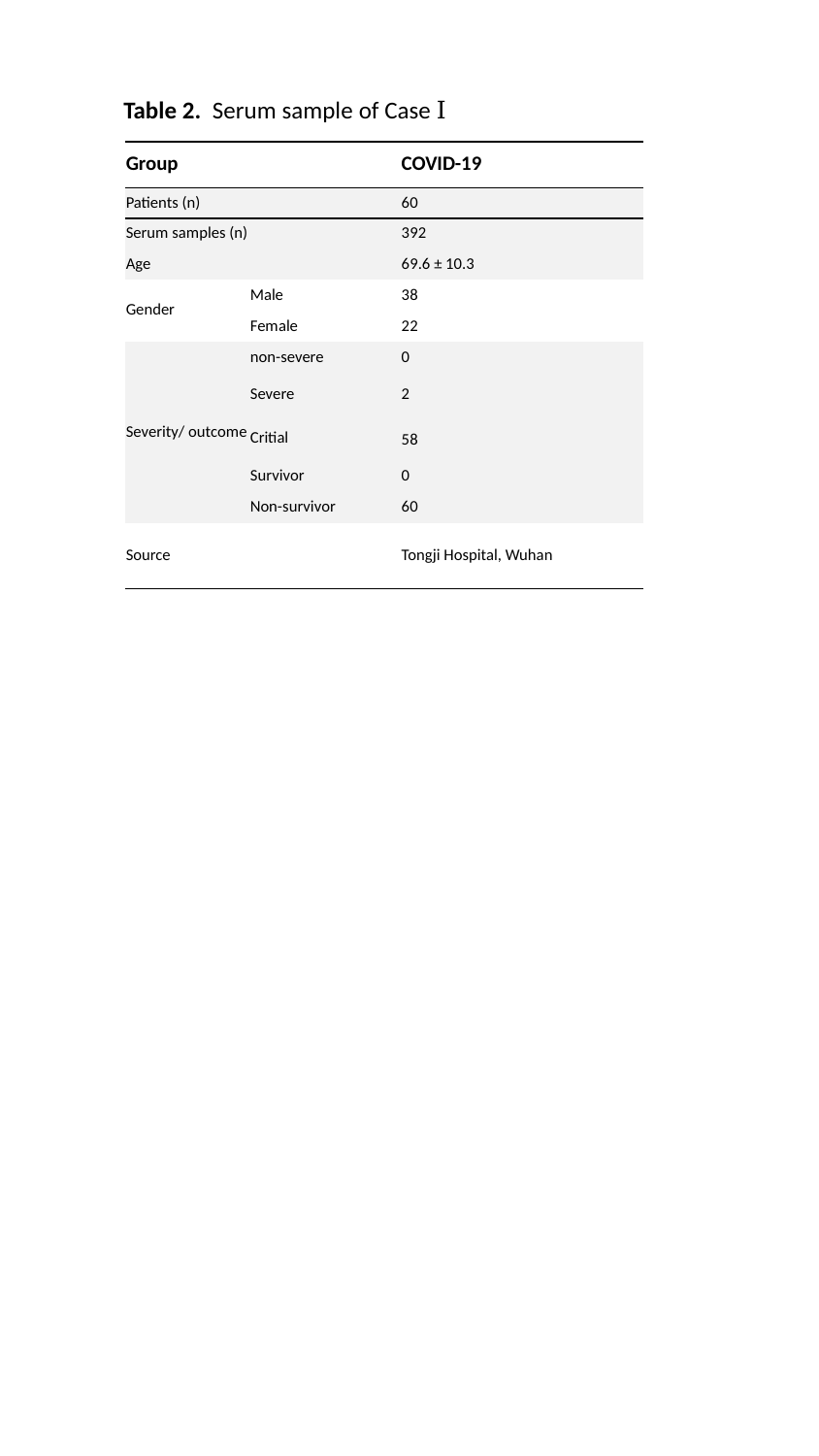

Table 2. Serum sample of Case Ⅰ
| Group | | COVID-19 |
| --- | --- | --- |
| Patients (n) | | 60 |
| Serum samples (n) | | 392 |
| Age | | 69.6 ± 10.3 |
| Gender | Male | 38 |
| | Female | 22 |
| Severity/ outcome | non-severe | 0 |
| | Severe | 2 |
| | Critial | 58 |
| | Survivor | 0 |
| | Non-survivor | 60 |
| Source | | Tongji Hospital, Wuhan |

### Slide 3
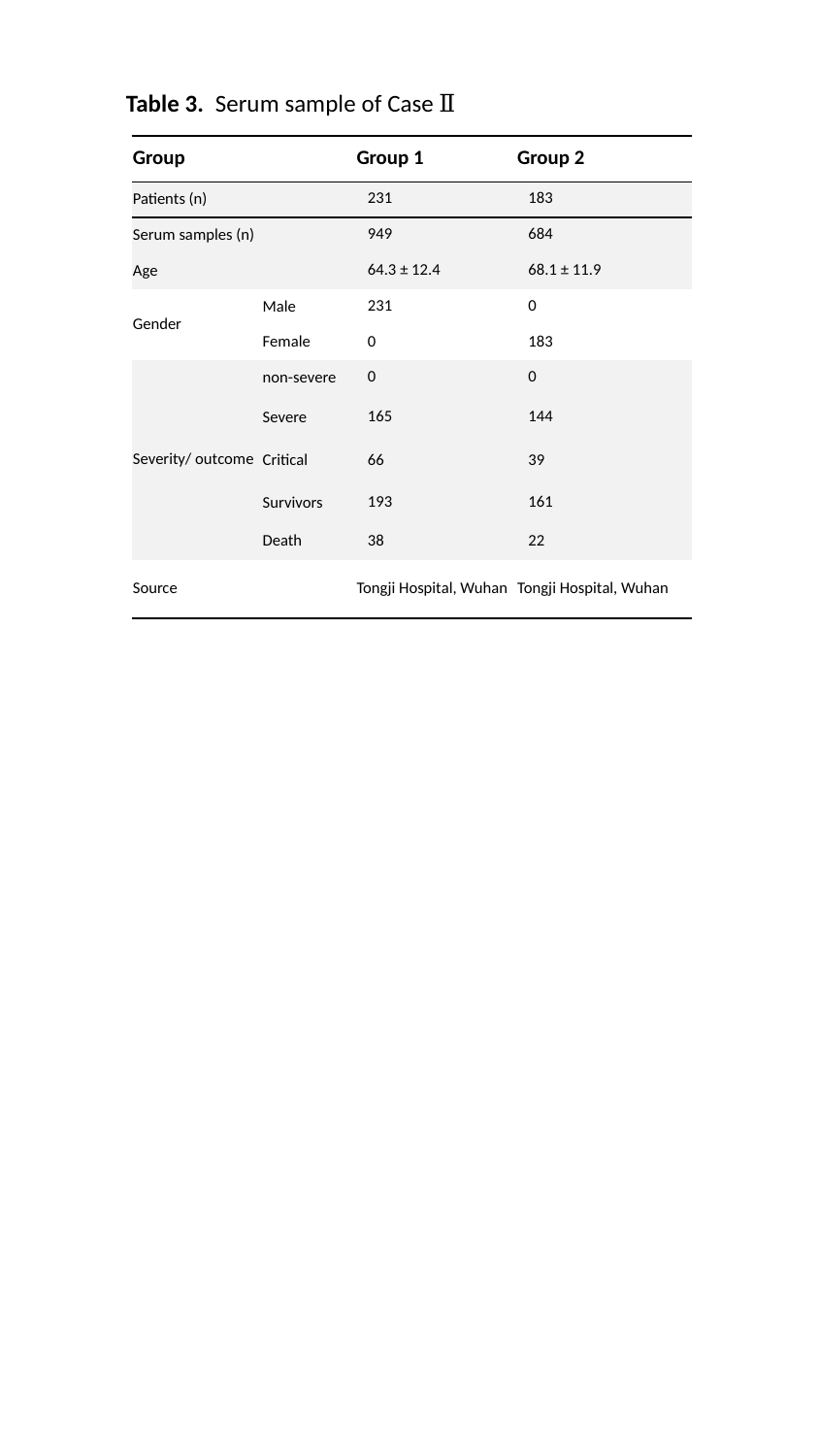

Table 3. Serum sample of Case Ⅱ
| Group | | Group 1 | Group 2 |
| --- | --- | --- | --- |
| Patients (n) | | 231 | 183 |
| Serum samples (n) | | 949 | 684 |
| Age | | 64.3 ± 12.4 | 68.1 ± 11.9 |
| Gender | Male | 231 | 0 |
| | Female | 0 | 183 |
| Severity/ outcome | non-severe | 0 | 0 |
| | Severe | 165 | 144 |
| | Critical | 66 | 39 |
| | Survivors | 193 | 161 |
| | Death | 38 | 22 |
| Source | | Tongji Hospital, Wuhan | Tongji Hospital, Wuhan |

### Slide 4
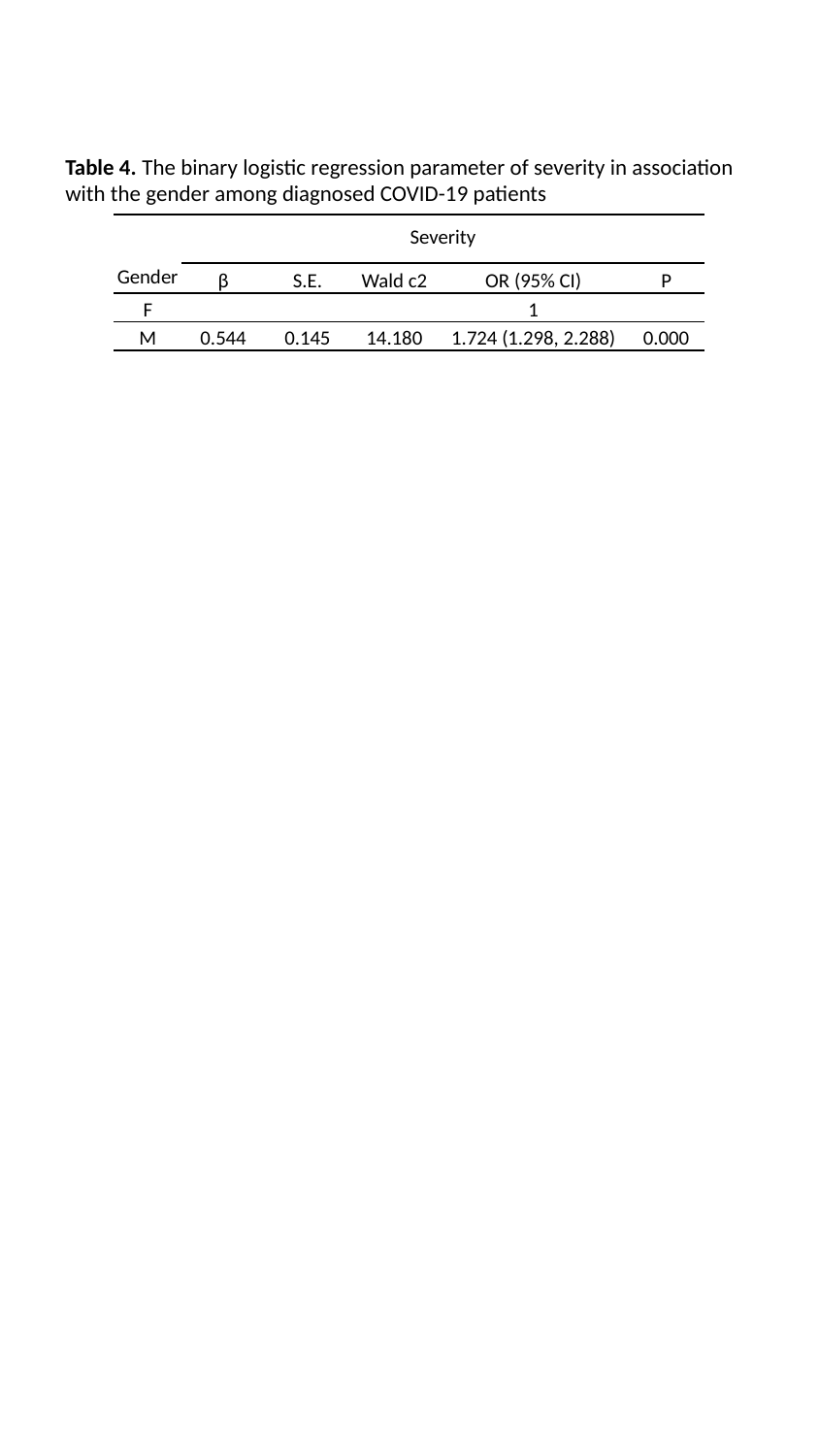

Table 4. The binary logistic regression parameter of severity in association with the gender among diagnosed COVID-19 patients
| | Severity | | | | |
| --- | --- | --- | --- | --- | --- |
| Gender | β | S.E. | Wald c2 | OR (95% CI) | P |
| F | | | | 1 | |
| M | 0.544 | 0.145 | 14.180 | 1.724 (1.298, 2.288) | 0.000 |
