## Supplementary file for "COVID-ONE-humoral immune: The One-stop Database for COVID-19-specific Antibody Responses and Clinical Parameters"

### Slide 1
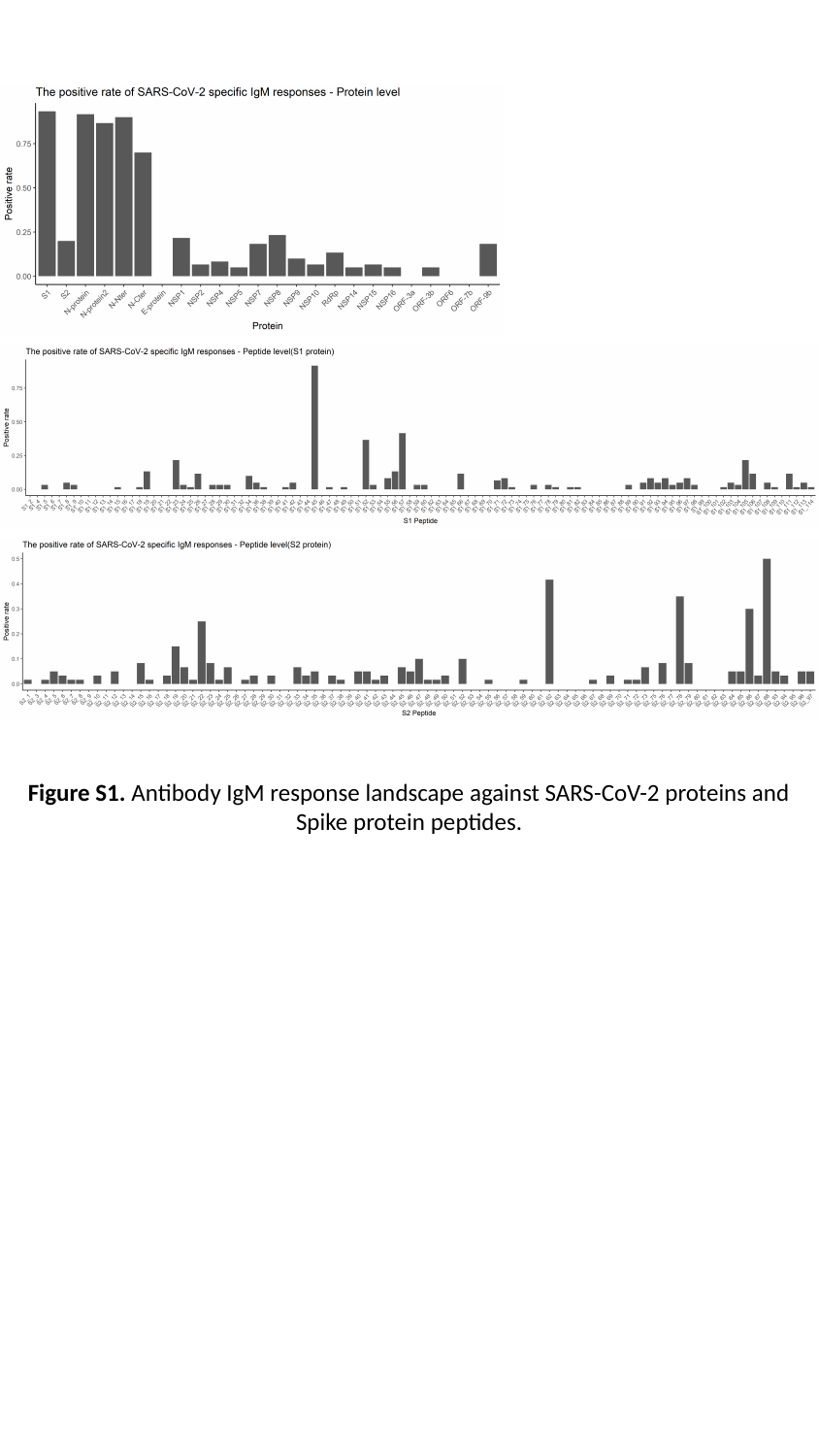

Figure S1. Antibody IgM response landscape against SARS-CoV-2 proteins and Spike protein peptides.

### Slide 2
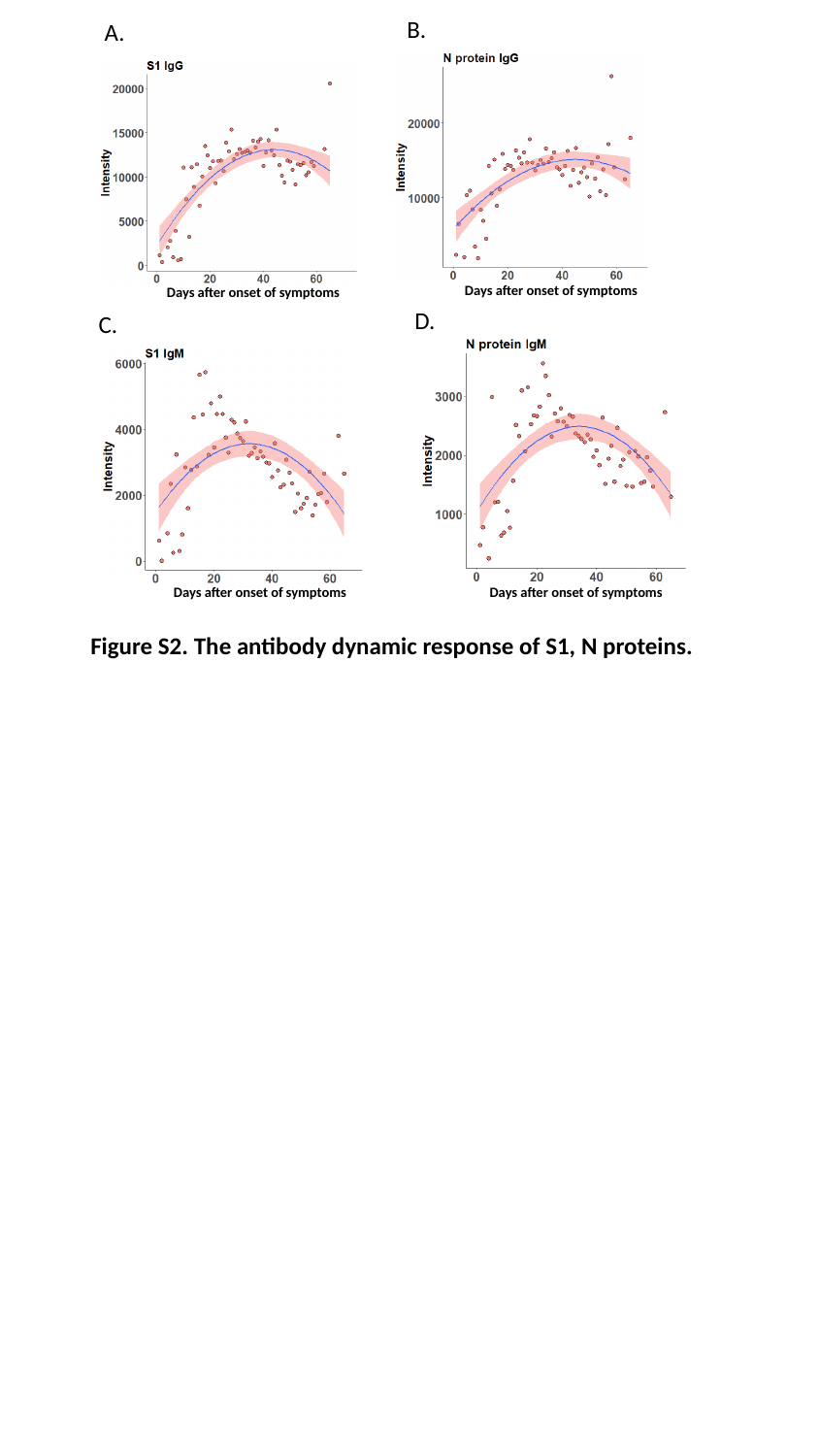

B.
A.
Days after onset of symptoms
Days after onset of symptoms
D.
C.
Days after onset of symptoms
Days after onset of symptoms
Figure S2. The antibody dynamic response of S1, N proteins.

### Slide 3
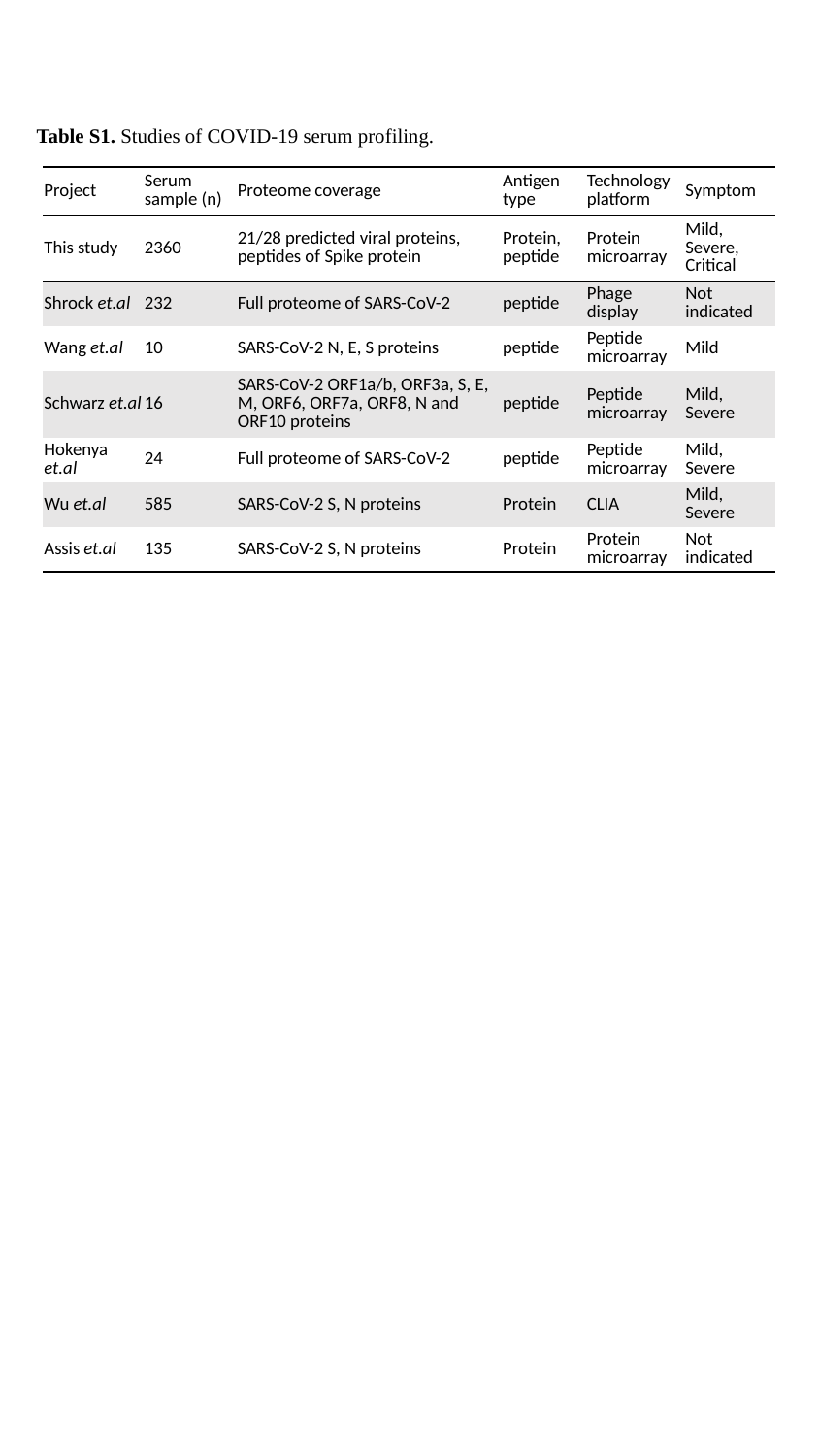

Table S1. Studies of COVID-19 serum profiling.
| Project | Serum  sample (n) | Proteome coverage | Antigen type | Technology platform | Symptom |
| --- | --- | --- | --- | --- | --- |
| This study | 2360 | 21/28 predicted viral proteins, peptides of Spike protein | Protein, peptide | Protein microarray | Mild, Severe, Critical |
| Shrock et.al | 232 | Full proteome of SARS-CoV-2 | peptide | Phage display | Not indicated |
| Wang et.al | 10 | SARS-CoV-2 N, E, S proteins | peptide | Peptide microarray | Mild |
| Schwarz et.al | 16 | SARS-CoV-2 ORF1a/b, ORF3a, S, E, M, ORF6, ORF7a, ORF8, N and ORF10 proteins | peptide | Peptide microarray | Mild, Severe |
| Hokenya et.al | 24 | Full proteome of SARS-CoV-2 | peptide | Peptide microarray | Mild, Severe |
| Wu et.al | 585 | SARS-CoV-2 S, N proteins | Protein | CLIA | Mild, Severe |
| Assis et.al | 135 | SARS-CoV-2 S, N proteins | Protein | Protein microarray | Not indicated |
